# Sequencing analysis of *Helicobacter pylori* infection in gastric mucosa and its progression to gastric mucosa-associated tissue (MALT) lymphoma

**DOI:** 10.1101/2024.04.12.589001

**Authors:** Seung-Joo Yoo, Youngil Koh, Ja Min Byun, Junshik Hong, Dong-Yeop Shin, Jaeyoung Chun, Hyunsoo Chung, Sung-Soo Yoon

**Author notes:** Co-corresponding authors, **Corresponding authors**, Jaeyoung Chun, MD, PhD, Division of Gastroenterology, Department of Internal Medicine, Yonsei University College of Medicine 20, Eonju-ro 63-gil, Gangnam-gu, Seoul, Republic of Korea, Seoul 06229, Korea, Hyunsoo Chung, MD, PhD, Department of Internal Medicine, Seoul National University College of Medicine 101, Daehak-ro, Jongno-gu, Seoul 03080, Republic of Korea, Sung-Soo Yoon, MD, PhD, Department of Internal Medicine, Seoul National University Hospital 101, Daehak-ro, Jongro-gu, Seoul 03080, Republic of Korea. Co-first authors (Seung-Joo Yoo, Youngil Koh).

## Abstract

**Introduction:** The pathogenesis of gastric mucosa-associated tissue (MALT) lymphoma is associated with *Helicobacter pylori* infection. Although treatment strategies and responsiveness according to the stage of gastric MALT lymphoma have been widely reported, a detailed study of the biological carcinogenic process is still required.

**Method:** Paired, fresh tumor-adjacent normal and gastric mucosal tissue samples from 13 patients with gastric MALT lymphoma were prospectively collected. Whole exome sequencing (WES) and whole-transcriptome sequencing (WTS) data were generated. The analysis of mutations, gene fusion, gene expression, and the microbiome was stratified by *H. pylori* infection and disease status.

**Results:** Somatic mutations in *TRAF3* and *TNFAIP3* were identified in *H. pylori*-negative gastric MALT lymphoma. Fusions involving *BIRC3-MALT1* (2 samples) and *TBL1XR1-TP63* (1 sample with *H. pylori* infection) were detected. Stepwise comparative analysis of RNA expression revealed upregulation of immune response, inflammatory responses, and the NF- κB signaling pathway in *H. pylori*-positive MALT lymphoma cases. Pathways associated with pathogens were upregulated in *H. pylori*-negative MALT lymphoma cases, suggesting that infections other than *H. pylori* may affect lymphomagenesis. Microbiome analysis revealed that *genus_Rothia* was negatively correlated with alpha diversity.

**Conclusion:** A stepwise approach using diverse stages of WTS data revealed detailed pathogenic mechanisms of gastric MALT lymphoma. Chronic inflammation following infection contributes to gastric MALT lymphomagenesis in both *H. pylori* positive and negative cases.

## Background

Gastric mucosa-associated lymphoid tissue (MALT) lymphoma is a low-grade non-Hodgkin lymphoma that originates from the post-germinal center B cells[1]. Gastric MALT lymphoma is the most common primary gastric lymphoma[2]. *Helicobacter pylori* (*H. pylori*) is associated with gastric MALT lymphoma; approximately 90% of patients with gastric MALT lymphoma are infected with *H. pylori* and 60–92% of cases respond to eradication therapy[3, 4].

In patients infected with *H. pylori*, the stomach is exposed to a low-pH environment and chronic stimulation, leading to the development of chronic gastritis and the acquisition of MALT[1, 5]. The development of MALT lymphoma is induced by genetic alterations and/or proliferation of monoclonal B cells following stimulation of *H. pylori*[5]. Translocations of t(1;2)(p22;p12)/*IGK*-*BCL10* and genetic alterations in *TNFAIP3*, *CARD11*, *among other things*, that indicate an abnormal activation of the NF-κB signaling pathway, play a fundamental role in driving gastric MALT lymphoma in both *H. pylori*-positive and *H. pylori*-negative cases[6].

In the early stages, MALT lymphoma depends on *H. pylori*, thus it is expected to regress with eradication therapy. However, in the advanced stages of the disease, MALT lymphoma transforms into an *H. pylori*-independent cancer, exhibiting a poor response to eradication therapy[7, 8]. The treatment strategies and responsiveness according to the stage of *H. pylori*- positive gastric MALT lymphoma have been reported[9–11]. However, detailed biological studies of this carcinogenic process are required. Although some studies have reported the results of microarray analysis of gastric MALT lymphoma, RNA-seq analysis using diverse stages of gastric mucosa samples would be helpful in elucidating the carcinogenesis of gastric MALT lymphoma by *H. pylori*[12–14].

In addition, in cases of *H. pylori*-negative gastric MALT lymphoma, infection with other microbiota, autoimmune diseases, and genetic alterations are believed to contribute to carcinogenesis[15]. The most common genetic alteration is the translocation of t(11;18)(q21;q21)/*BIRC3*-*MALT1*, which is also associated with poor response to eradication therapy in *H. pylori*-negative cases[8, 16]. Interestingly, some patients without translocations respond to eradication even though they were not infected with *H. pylori*[17]. Considering the pathophysiology of *H. pylori*-positive cases and their phenotypic similarity to *H. pylori*- negative cases, a microbiome other than *H. pylori* may contribute to the development of *H. pylori*-negative gastric MALT lymphoma. Similarly, the differences in the microbiome between *H. pylori*-negative gastric MALT lymphoma and control patients have also been reported[18]. In summary, these findings highlight the importance of analyzing the microbiota of *H. pylori*-negative gastric MALT lymphomas.

Therefore, the current study aimed to elucidate the pathways that contribute to the carcinogenesis of MALT lymphoma following microbial infection. Additionally, we aimed to identify the microbiome responsible for initiating carcinogenesis in *H. pylori*-negative cases. For this purpose, the genetic changes and RNA expression patterns of gastric MALT lymphoma, adjacent normal tissue, and follow-up gastric mucosal tissue were compared by prospectively collecting fresh samples from patients with newly diagnosed gastric MALT lymphoma.

## Methods

### Patients and samples

A total of thirteen patients diagnosed with gastric MALT lymphoma were included in this study. Paired fresh samples of tumors and adjacent normal gastric mucosal tissues were collected using endoscopy before treatment. Additionally, follow-up gastric mucosal tissue and saliva samples were collected. To identify whether patients were infected with *H. pylori*, we conducted an Immunoglobulin G test, *Campylobacter*-like organism test, Urea Breath Test, and pathological tests on patients and samples. This study was approved by the IRB (Institutional Review Board) of Seoul National University Hospital (No. H-1704-070-845).

### Sequencing data generation

The DNA collected from the sample was used for whole exome sequencing (WES) and whole transcriptome sequencing (WTS). Library construction for WES data generation was performed using the SureSelect XT Human All Exon Kit V5 (Agilent, CA, USA), followed by sequencing on an Illumina HiSeq2500 platform with a 2 × 100 read length. To generate the WTS data, libraries were prepared using the TruSeq Stranded Total RNA Kit with Ribo-Zero Globin (Illumina, San Diego, CA). Paired-end sequencing of these libraries was conducted using the Illumina HiSeq2500 (Illumina, San Diego, CA) in accordance with the manufacturer’s instructions. Of the 13 cases, only 10 sets of WES and WTS data were generated; three patients provided only WTS data.

### Somatic mutation calling & filtering

To identify somatic mutations in cancer samples, whole-exome sequencing of cancer and saliva DNA samples was used. The quality of the sequenced reads was initially assessed using Fastqc (v.0.11.9)[19]. Low-quality reads were removed using Trimmomatic (version 1.39)[20]. The filtered sequencing reads were aligned to the human reference genome (GRCh37/hg19) using the Burrows–Wheeler Aligner[21]. Following alignment, somatic SNVs (Single Nucleotide Variations)/indels were identified using the Genome Analysis Tool Kit (GATK, v.4.2.0.0; https://software.broadinstitute.org/gatk/)[22]. The somatic mutations were identified using Mutect2 of GATK and the biological information was annotated using AnnoVar and VEP[23, 24]. Using these annotation tools, information from the 1000 Genomes Project, gnomAD, ClinVar, and Catalogue of Somatic Mutations in Cancers (COSMIC, version 96)[25–28] was added. To filter out sequencing artifacts, variants with a depth lower than 50 and variants with allelic fractions between 7% and 40% were excluded. Additionally, variants with strand bias between 0.25 and 4 were included. We removed variants reported in origins other than hematopoietic lymphoid tissue and stomach. In the gnomAD, variants with an allele frequency of 0.1 or more in the East Asian region were excluded.

### Subgroups of samples

The samples were divided into subgroups based on the identification of *the presence of H. pylori*. As mentioned previously, multiple tests were conducted on each sample. To judge the presence of *H. pylori*, an IgG test was prioritized, and other tests supported this decision.

### Differentially expressed genes & gene set enrichment analysis

The RNA expression levels were estimated using whole-transcriptome sequencing data. The sequenced reads were aligned to the human reference genome (GRCh37/hg19) using STAR (Spliced Transcript Alignment in Reference v.3.6)[29]. RNA expression normalization and read count calculations were performed using RSEM (RNA-seq by Expectation Maximization, v.1.3.2)[30]. Using the estimated RNA expression levels, differentially expressed genes (DEG) were analyzed across samples in multiple comparison groups and gene set enrichment analysis (GSEA) was performed using R software (version 4. 2. 2) and the R package, DESeq2 (v.1.38.3)[31, 32].

### Detection of genetic translocation

The Genetic translocations in the tumor samples were identified using three detection tools: Fusioncatcher, STAR-Fusion, and Arriba[33–35]. Raw FASTQ files were used as input files for Fusioncatcher and STAR-Fusion, whereas Arriba was fed with BAM files generated using STAR-Aligner processing[29]. To remove possible false-positive variants, only variants detected by all three tools were retained. Furthermore, fusions with limited supporting evidence reads were filtered out and assessed as false positives using an integrative genomics viewer[36].

### Detection & analysis of the mucosal microbiome

To identify the microbiome from the WTS data, a microbiome detection protocol was followed using Kraken software[37, 38]. Before classifying the microbiome, it was isolated from the transcriptome data by excluding reads mapped to the human reference genome (GRCh37/hg19). Subsequently, the reads were classified by mapping them to a reference sequence of the microbiota. During the process, the minimum hit groups were set to three, as recommended in the protocol, and a confidence score of 0.10 to filter out the reads with insufficient supporting evidence[37].

## Results

The clinical characteristics of the 13 patients are shown in Table 1. Based on *H. pylori* status, location, and timing of the samples, the samples were categorized into six subgroups. As a result, 37 WTS data points from 13 patients with gastric MALT lymphoma were classified into *H. pylori*-positive cancer, *H. pylori*-positive normal, *H. pylori*-negative cancer, *H. pylori*- negative normal, *H. pylori*-positive follow-up, and *H. pylori*-negative follow-up groups (Table 1).

**Table 1.** Clinicopathologic features of gastric MALT lymphoma. MB: mid body, LB: lower body, PW: post wall, GC: great curvature, AW: ant wall, LC: less curvature.

| No | Age | Gender | Location | *H. pylori* |
| --- | --- | --- | --- | --- |
| 1 | 69 | M | MBPW | positive |
| 2 | 37 | M | Angle | Positive |
| 3 | 37 | F | AntrumGC | Negative |
| 4 | 57 | M | AntrumGC | Negative |
| 5 | 71 | F | AntrumAW | Positive |
| 6 | 68 | F | MBPW | Negative |
| 7 | 67 | F | AntrumGC | Negative |
| 8 | 47 | F | MBGC | Positive |
| 9 | 71 | F | LBPW | Positive |
| 10 | 53 | F | LBLC | Positive |
| 11 | 52 | F | AntrumGC | Positive |
| 12 | 60 | M | MBGC | Positive |
| 13 | 57 | F | LBAW | Positive |

### Genetic translocation

To identify recurrent or novel gene fusions in gastric MALT lymphoma, genetic translocations from the WTS data were detected using Fusioncatcher, STAR-Fusion, and Arriba. We identified two genetic translocations: t(11;18)(q21;q21)/*BIRC3*-*MALT1* and inv(3)(q26q28)/*TBL1XR1*-*TP63* (Figure 1B). *BIRC3-MALT1* gene fusions were identified in *H. pylori*-negative samples, consistent with previous reports on recurrent cases of *H.* pylori-negative gastric MALT lymphoma patients[39]. Additionally, an inversion between *TBL1XR1*–*TP63* was found in *H. pylori*-positive samples. The *TBL1XR1*–*TP63* gene fusion has been reported in a study of transcriptome data of diffuse large B-cell lymphoma cases[40].

**Figure 1 A-B.**
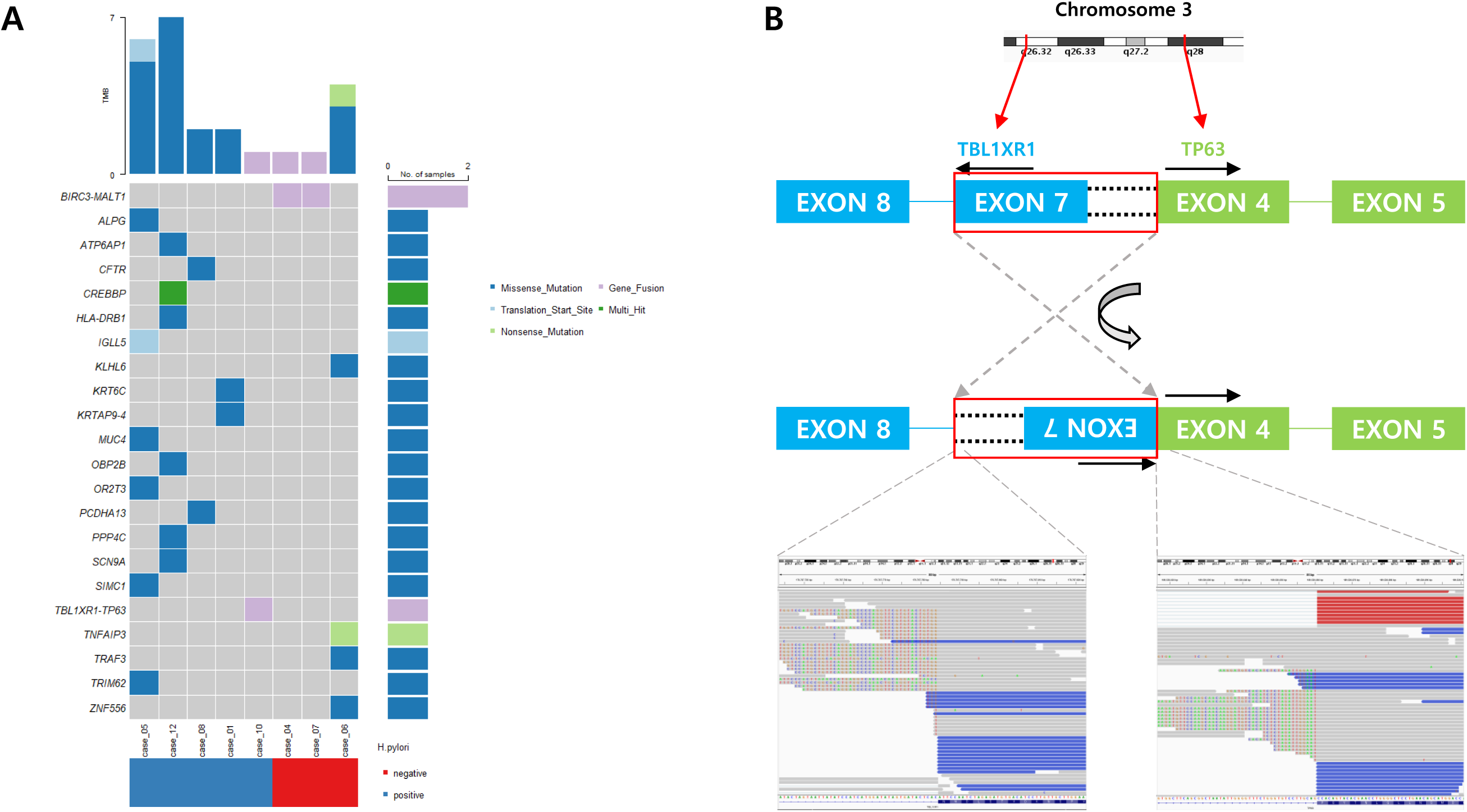
**A.** An oncoplot represents the somatic mutations from WES data including genetic translocations from WTS data. **B.** A schematic diagram represents the inversion of *TBL1XR1* and *TP63*. Inversion of *TBL1XR1* and *TP63.* A schematic diagram shows detailed breakpoints of the genetic translocations. The screenshot of IGV shows the evidence sequencing reads from the cancer sample.

### Somatic mutation

From the 10 WES data, somatic mutations were identified and false-positive mutations were filtered out. Twenty gene mutations were identified in five samples (two *H. pylori*-positive and three *H. pylori*-negative samples). *TNFAIP3*, *TRAF3*, and *KLHL6*, which were mutated in *H. pylori*-negative gastric MALT lymphoma cases, were also mutated in *H. pylori*-negative samples (case 06). No somatic mutations were found in the cases (case 04 and case 07) with genetic translocations, t(11;18)(q21;q21)/*BIRC3*-*MALT1* (Figure 1A). *MUC4*, *TRIM*, *ALPG*, and other genes found in other cancer types were detected in both *H. pylori*-negative and *H. pylori*-positive cancer samples.

### Gene expression analysis

To analyze the differentially expressed genes among the sample subgroups, we conducted multiple statistical tests using R statistical software. Our objective was to understand the transcriptional differences between the following samples at various stages of gastric MALT lymphoma carcinogenesis: 1) *H. pylori*-positive normal samples versus *H. pylori*-negative normal samples; 2) *H. pylori*-positive cancer samples versus *H. pylori*-positive normal samples; 3) *H. pylori*-positive cancer samples versus *H. pylori*-negative cancer samples; and 4) *H. pylori*-negative cancer samples & *H. pylori*-negative normal samples.

#### 1) *H. pylori*-positive normal samples & *H. pylori*-negative normal samples

The differences in gene expression between *H. pylori*-positive and *H. pylori*-negative normal samples were investigated to identify biological changes in the gastric mucosa when infected with *H. pylori*. Eleven down-regulated and two upregulated genes were statistically significant in the *H. pylori*-positive group. Of these, *MUC4*, *CXCL5*, *RNU4-2*, *NLRP7* and *IGKV2-24*, associated with regulating pH in the gastric environment and the proton pump, were down- regulated (Figure 2B; Table 2). From the results of the GSEA using Gene Ontology (GO) terms, the response to bacteria, immune response-regulating signaling pathway, inflammatory response, T cell activation, and B cell activation were upregulated in *H. pylori*-positive normal samples. TNF-α signaling and interferon gamma response were upregulated in *H. pylori*- positive normal samples, which was similar to the study that reported increased levels of cytokine, interferon-γ, and tumor necrosis factor-α in the *H. pylori*-infected human stomach [41]. Furthermore, upregulated B cell-mediated immunity and B cell receptor signaling pathways were found, which could develop into the oncogenic pathway of gastric MALT lymphoma.

**Figure 2 A-C.**
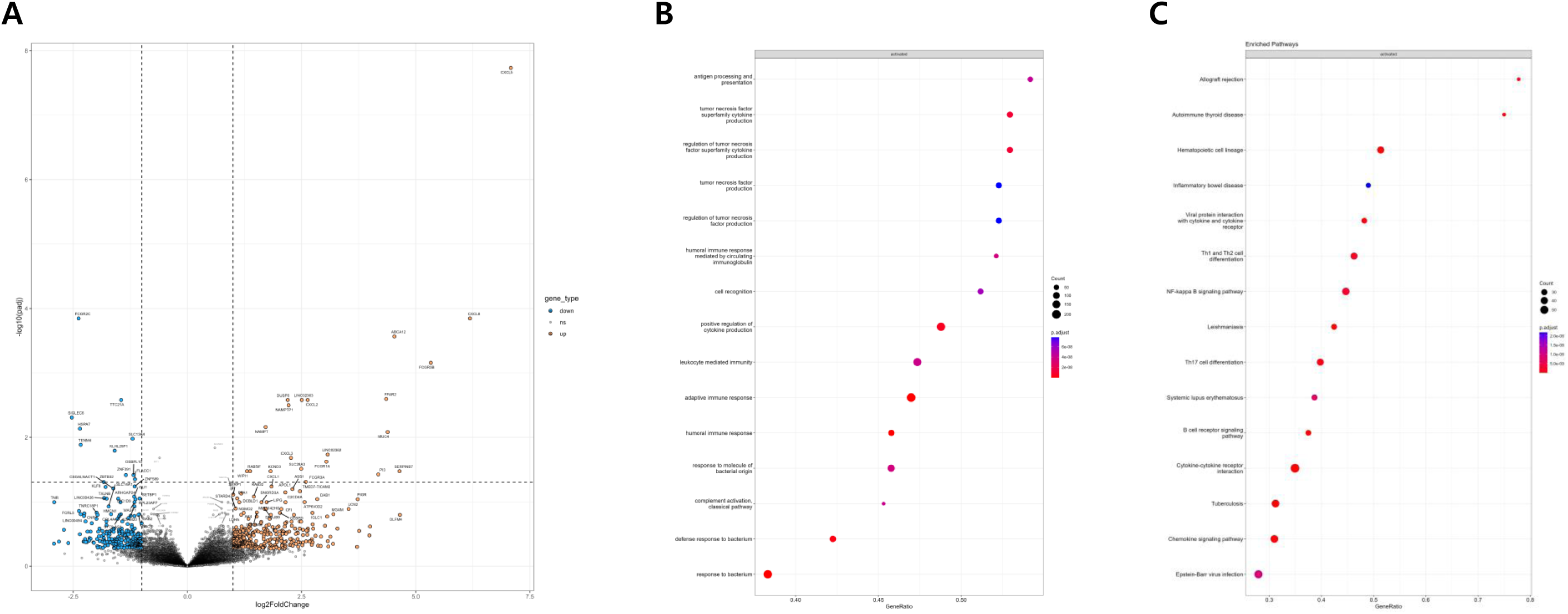
**A. A** volcano plot shows the results of DEG analysis between *H. pylori*-positive cancer samples and *H. pylori*-negative cancer samples. Genes that have the following range of fold changes are marked with blue and red colors and the dashed line shows a cut-off of adjusted p-value and fold changes. **B-C.** The two plots show the results of GSEA from “*H. pylori*-positive normal samples & *H. pylori*-negative normal samples” and “*H. pylori*-negative cancer samples & *H. pylori*-negative normal samples.”

**Table 2.** A table displays the results of DEG analysis from multiple comparisons. The genes in this table were selected by log2FoldChange and padj values and the full set of genes which were statistically significant are listed in the supplemental Table S1-.

| GENE | log2FoldChange | padj |
| --- | --- | --- |
| *H. pylori*-positive normal vs *H. pylori*-negative normal |  |  |
| *MUC4* | 4.768 | 1.900E-17 |
| *CXCL5* | 6.539 | 2.089E-09 |
| *RNU4-2* | -3.316 | 2.179E-05 |
| *NLRP7* | 4.434 | 3.245E-04 |
| *IGKV2-24* | 3.369 | 6.311E-04 |
| *H. pylori*-positive cancer vs *H. pylori*-positive normal |  |  |
| *SELL* | 2.241 | 1.035E-06 |
| *HLA-DMB* | 1.241 | 2.269E-06 |
| *NKX6-3* | -4.902 | 5.988E-06 |
| *IRF8* | 1.189 | 3.295E-05 |
| *TMEM178B* | 1.453 | 3.476E-05 |
| *EIF3CL* | 7.380 | 3.869E-05 |
| *CXCR4* | 1.577 | 3.878E-05 |
| *H. pylori*-positive cancer vs *H. pylori*-negative cancer |  |  |
| *CXCL5* | 7.085 | 1.843E-08 |
| *CXCL8* | 6.188 | 1.425E-04 |
| *FCGR2C* | -2.382 | 1.425E-04 |
| *ABCA12* | 4.532 | 2.719E-04 |
| *FCGR3B* | 5.330 | 6.975E-04 |
| *FFAR2* | 4.352 | 2.534E-03 |
| *CXCL2* | 2.633 | 2.631E-03 |
| *H. pylori*-negative cancer vs *H. pylori*-negative normal |  |  |
| *CNR1* | 4.750 | 2.195E-06 |
| *RUBCNL* | 4.350 | 6.220E-06 |
| *C3* | 4.167 | 6.220E-06 |
| *AIM2* | 4.766 | 6.220E-06 |
| *CD37* | 4.089 | 6.431E-06 |
| *CCL19* | 4.952 | 6.430E-06 |

#### 2) Comparison of *H. pylori*-positive cancer & *H. pylori*-positive normal samples

When comparing *H. pylori*-positive cancer and *H. pylori*-positive normal samples, we anticipated discovering oncogenic dysregulation of genes and pathways, such as the NF-κB and the B cell signaling pathways, as previously reported[42, 43].

From the DEG analysis, 263 upregulated and 128 down-regulated genes were observed. Of the significantly dysregulated genes in *H. pylori*-positive cancer samples, *SELL*, *HLA-DMB*, *CXCR4* and *EIF3CL* were upregulated, whereas *NKX6-3*, *GAST*, and *MEIS1* were down- regulated (Table 2). Following GSEA, upregulated immune and inflammatory responses in *H. pylori*-positive cancer samples were observed. Upregulation of the NF-κB signaling pathway, B cell proliferation, B cell signaling pathway via JAK-STAT, and hematopoietic cell lineage in cancer samples were found, all of which are considered as oncogenic pathways. These results are consistent with those of a previous study showing that dysregulation of oncogenic pathways following chronic inflammation induced by *H. pylori* plays a significant role in the development of gastric MALT lymphoma[2, 5, 6].

#### 3) *H. pylori*-positive & *H. pylori*-negative cancer samples

The analysis of DEGs revealed 22 upregulated and 12 down-regulated genes between *H. pylori*-positive and *H. pylori*-negative cancer samples. In that set, CXCL5, CXCL8 and CXCL2 were up-regulated (Figure 2A). Gene set enrichment analysis indicated that immune response, inflammatory response, and pathogen infection were upregulated in the *H. pylori*- positive cancer group; this supports inflammation-related carcinogenesis in *H. pylori*-infected cases. The absence of a difference in the carcinogenic pathways, such as the NF-κB signaling pathway, suggests common carcinogenesis at the molecular level regardless of *H. pylori* infection status. Moreover, the causes of *H. pylori*-negative gastric MALT lymphoma may include gene fusion events, infections with other microbiota, and autoimmune diseases. Of the *H. pylori*-negative samples, two exhibited the t(11;18)(q21;q21)/*BIRC3*-*MALT1* gene fusion. None of the samples showed characteristics of an autoimmune disease.

#### 4) *H. pylori*-negative cancer & *H. pylori*-negative normal samples

To investigate the potential drivers of *H. pylori*-negative gastric MALT lymphoma, DEG and GSEA were performed on *H. pylori*-negative cancer and *H. pylori*-negative normal samples. A total of 766 genes were differentially expressed between the groups. The upregulated genes in cancer tissues included *CNR1, RUBCNL*, *C3, AIM2, CD37* and *CCL19,* whereas *C11orf86*, *EDN3*, *CHGB*, and *SST* were down-regulated in cancer samples (Table 2). GSEA revealed that the GO terms (B cell receptor signaling pathway, B cell proliferation, and immune responses) and the KEGG (NF-κB and JAK-STAT signaling pathways) were upregulated in cancer samples (Figure 2C). Additionally, the activation of pathways related to *Staphylococcus aureus* infection, tuberculosis, and toxoplasmosis in KEGG terms was detected, suggesting that some of the *H. pylori*-negative samples may be related to infections other than *H. pylori*.

### Microbiome analysis

Using kraken2, a k-mer-based taxonomic classification, sequencing reads not aligned to the human reference genome, including follow-up samples, were classified. A total of 37 WTS data points were used to detect microbiomes. Initially, the relative abundance of the microbiome at the genus level was analyzed. Some cases that were *H. pylori*-positive had no *g_Helicobacter*, whereas some *H. pylori*-negative cases had *g_Helicobacter*. Across all samples, various microbiomes were detected, but certain genera consistently exhibited high abundance: *g_Helicobacter, g_Rothia, g_Veillonella*, and *g_Porphyromonas* (Figure 3A).

**Figure 3.**
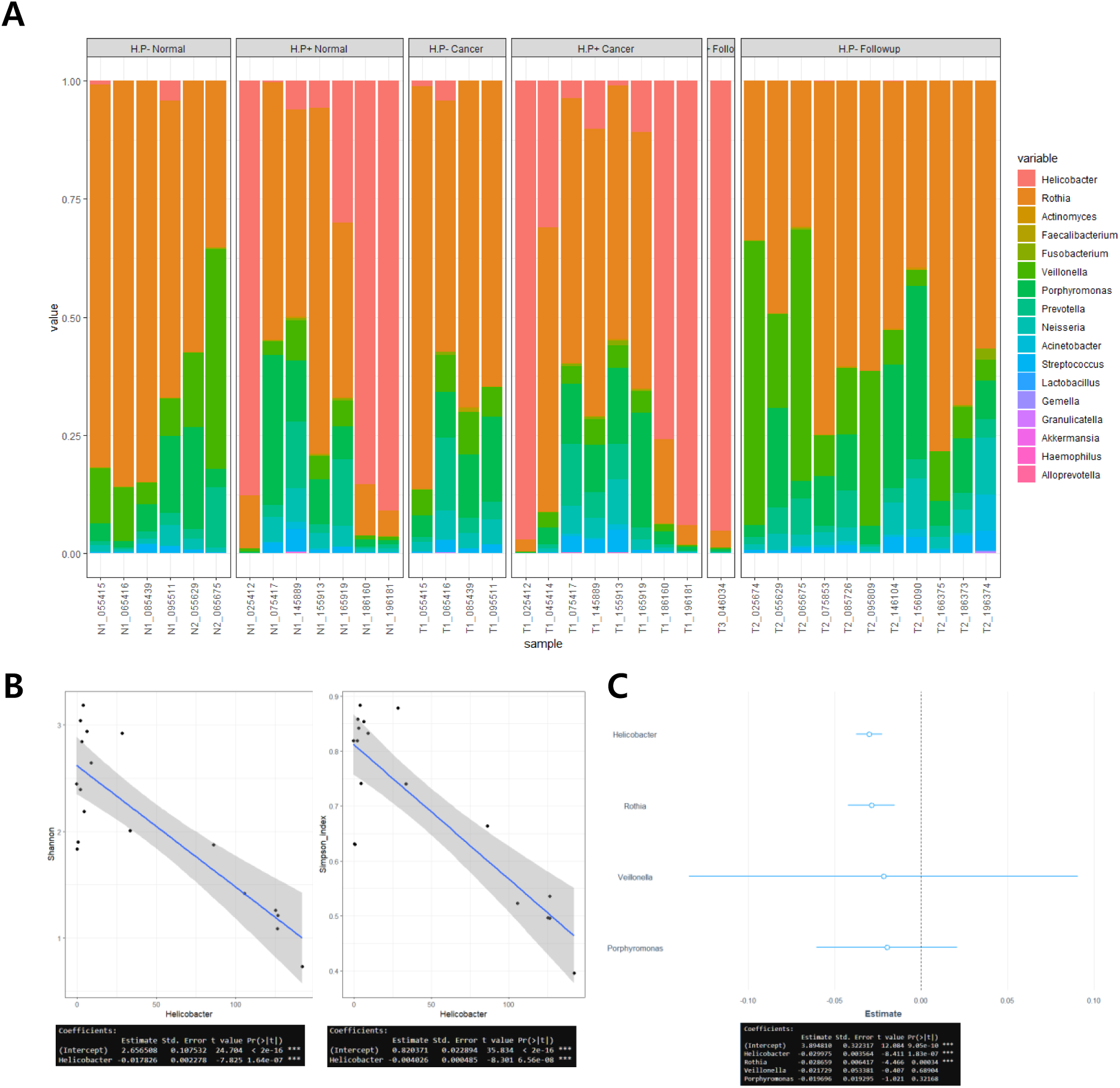
**A-C** : A. A stacked bar shows a relative abundance of the microbiome from all WTS data. The designation of each group means the presence of *H. pylori* and tissue type. B. Each dot represents an index value matching the quantified read count of *H. pylori*. C. The dots and lines represent the estimate and error term from multivariate linear regression.

As previously reported, patients infected with *H. pylori* have a lower alpha diversity than *H. pylori*-negative cases[44]. The alpha diversities of the samples were investigated using the Shannon and Simpson indices, and no difference between *H. pylori*-positive and *H. pylori*- negative samples was found (p = 0.4). Therefore, a linear regression analysis was conducted to determine whether *H. pylori* was associated with alpha diversity, and to assess the impact of other microbiomes on alpha diversity. The quantified counts of the sequenced microbiome reads were used as independent variables and alpha diversity was used as the response variable.

In the univariate linear regression, among the four genera (*g_Helicobacter, g_Rothia, g_Veillonella*, and *g_Porphyromonas*) only *g_Helicobacter* was correlated with alpha diversity (Shannon: beta= 0.017826, p = 1.64×10^-7^ / Simpson: beta= -0.004026, p = 6.56×10^-8^).

As alpha diversity decreased with an increase in *g_Helicobacter* reads, a multivariate linear regression with the four genera was conducted to identify linearly-related and independent variables. Both *g_Rothia* and *g_Helicobacter* were negatively correlated with alpha diversity (Figure 3B–C). In general, low alpha diversity is associated with unfavorable gut microbiome conditions related to chronic illnesses. Considering the correlation between *g_Helicobacter* and *g_Rothia*, and alpha diversity, we suggest that *g_Rothia* may be a potential factor contributing to *H. pylori*-negative gastric MALT lymphoma.

## Discussion

In this study, we prospectively collected samples and generated next-generation sequencing (NGS) data from patients with gastric MALT lymphoma and divided them into two groups for further analysis: *H. pylori*-positive and *H. pylori*-negative. WES and WTS data were produced and utilized to analyze genetic alteration, DEGs, GSEA, and microbiome detection.

Using fusion gene analysis, the *BIRC3*–*MALT1* gene fusion in *H. pylori*-negative cases was identified, consistent with previous findings[15–17]. Additionally, we discovered an inversion of *TBL1XR1*–*TP63* in *H. pylori*-positive cases. The *TBL1XR1–TP63* gene fusion has been reported as a recurrent somatic gene fusion in high-grade lymphomas, such as diffuse large B cell lymphoma (DLBCL) and is known to drive tumor survival through *EZH2* and upregulate *MYC*, *EZH2*, and *EED*[40, 45]. Indeed, the sample with the *TBL1XR1–TP63* gene fusion in the current study showed high expression of *MYC, EZH2,* and *EED* (supplemental Figure 1). In addition, patients with the *TBL1XR1*–*TP63* gene fusion showed a poor response to *H. pylori* eradication therapy. We speculate that the *TBL1XR1*–*TP63* fusion is one of the drivers of the high-grade transformation of gastric MALT lymphoma. Biological studies suggest B-cell lymphoma with *TBL1XR1–TP63* gene fusion might be susceptible to EZH1/2 inhibition[45].

Using RNA sequencing data from the fresh gastric mucosa of various clinical statuses, the biological phenomena associated with gastric MALT lymphomagenesis were successfully revealed. A multi-step, comparative analysis was conducted to detail the mechanism of gastric MALT lymphoma caused by *H. pylori*. First, by comparing *H. pylori*-positive and negative normal groups, the biological changes that occurred in the gastric mucosa upon infection with *H. pylori* were identified. It was expected that immune and inflammatory responses would be upregulated in the *H. pylori*-positive group due to the infection with *H. pylori*, which was indeed the case. The TNF-α signaling pathway was upregulated in the *H. pylori*-positive group, consistent with the studies that *H. pylori* induces TNF-α in the gastric mucosa[46, 47]. When comparing the *H. pylori*-positive cancer and *H. pylori*-positive normal groups, upregulation of oncogenic pathways, such as the NF-κB and B cell signaling pathways, was observed in the cancer group. Furthermore, in *H. pylori*-infected patients, the cancer group exhibited upregulation of chronic inflammation, immune response, and other pathways associated with the bacterial response in comparison with the normal group. The elevated expression of *CXCL5* and *CXCL8*, which had been studied as being upregulated by the inflammatory response after *H. pylori* infection, is also observed in the cancer group infected with *H. pylori*[48]. This suggests that inflammation triggered by *H. pylori* continues to occur in the cancer group. The upregulation of the immune response and other factors in *H. pylori*-positive compared to *H. pylori*-negative gastric MALT lymphoma further support this hypothesis. In summary, our stepwise approach confirmed that chronic inflammation and immune responses play critical roles in *H. pylori* lymphomagenesis. These findings suggest that for the treatment of gastric MALT lymphoma with *H. pylori* infection, immunotherapy agents may yield benefits.

It is also worth mentioning the pathophysiology of *H. pylori*-negative gastric MALT lymphomas using the stepwise approach. In the comparison of cancer and normal samples within the *H. pylori*-negative cases, aberrant NF-κB and B cell signaling pathways were dominant in lymphoma samples. This concurs with the studies that identified the upregulated NF-κB signaling pathway induced by some recurrent translocations [t(1;14)(p22;q32), t(14;18)(q32;q21), and t(11;18)(q21;q21)] in MALT lymphoma[49, 50]. Furthermore, upregulated biological pathways associated with pathogen infection were identified in the *H. pylori*-negative cancer compared to the *H. pylori*-negative normal group. These findings deepen our understanding of the molecular pathogenesis of *H. pylori*-negative gastric MALT lymphomas.

Based on the dysregulated pathways potentially influenced by specific microbiomes, we explored whether microbiomes other than *H. pylori* could be linked to gastric MALT lymphoma. There was a time when it was believed that no microorganisms could survive in the stomach; however, the discovery of *H. pylori* confirmed that microorganisms can thrive in the gastric environment. With advancements in NGS and analytical technologies, it has become evident that, in addition to *H. pylori*, various other microorganisms can interact within the gastric milieu. Studies have been conducted in Korean cohorts to analyze the stomach microbiome[51, 52]. Apart from the *g_Helicobacter*, diverse microbiota have been identified. In the current study, reads that were not mapped to the human reference genome in the WTS data were reclassified as part of the microbiome analysis, facilitating the detection of the microbiome present in each sample. Relative abundance and species diversity were investigated. The *g_Rothia* generally exhibited high abundance at the genus level, but several other genera also displayed noteworthy abundance. As observed in previous studies, *H. pylori*- infected patients exhibited lower alpha diversity at the genus level, as confirmed by linear regression. Moreover, multivariate linear regression suggested an association between *g_Rothia* and low alpha diversity, particularly in conjunction with *g_Helicobacter*. Low alpha diversity is considered indicative of an unhealthy microbial environment and has been studied in relation to chronic diseases. Although a direct association between low diversity and gastric MALT lymphoma has not been firmly established, given the confirmed association with *g_Helicobacter*, it is plausible that the *g_Rothia* plays a significant role in the gastric environment and may contribute to gastric MALT lymphoma.

## List of abbreviations

DEG: differentially expressed gene
GO: Gene Ontology
GSEA: gene set enrichment analysis
KEGG: Kyoto Encyclopedia of Genes and Genomes
MALT: gastric mucosa-associated tissue
NGS: next-generation sequencing
WES: whole exome sequencing
WTS: whole-transcriptome sequencing

## Declarations

Not applicable

## Ethics approval and consent to participate

All research was approved by the Institutional Review Board (IRB) of Seoul National University Hospital (IRB No. H-1704-070-845). All participants provided written informed consent to participate in this study.

## Consent for publication

Not applicable

## Availability of data and materials

Sequence data that support the findings of this study have been deposited in the Sequence Read Archive with the primary accession code PRJNA1088908.

## Authors’ contributions

## Acknowledgements

We thank the Global Science experimental Data hub Center (GSDC) and Korea Research Environment Open NETwork (KREONET) service for data computing and network provided by the Korea Institute of Science and Technology Information (KISTI).

## Authors’ information (optional)

Seung-Joo Yoo and Youngil Koh contributed equally to this work

