## Supplemental Figure 1 for "Sequencing analysis of *Helicobacter pylori* infection in gastric mucosa and its progression to gastric mucosa-associated tissue (MALT) lymphoma"

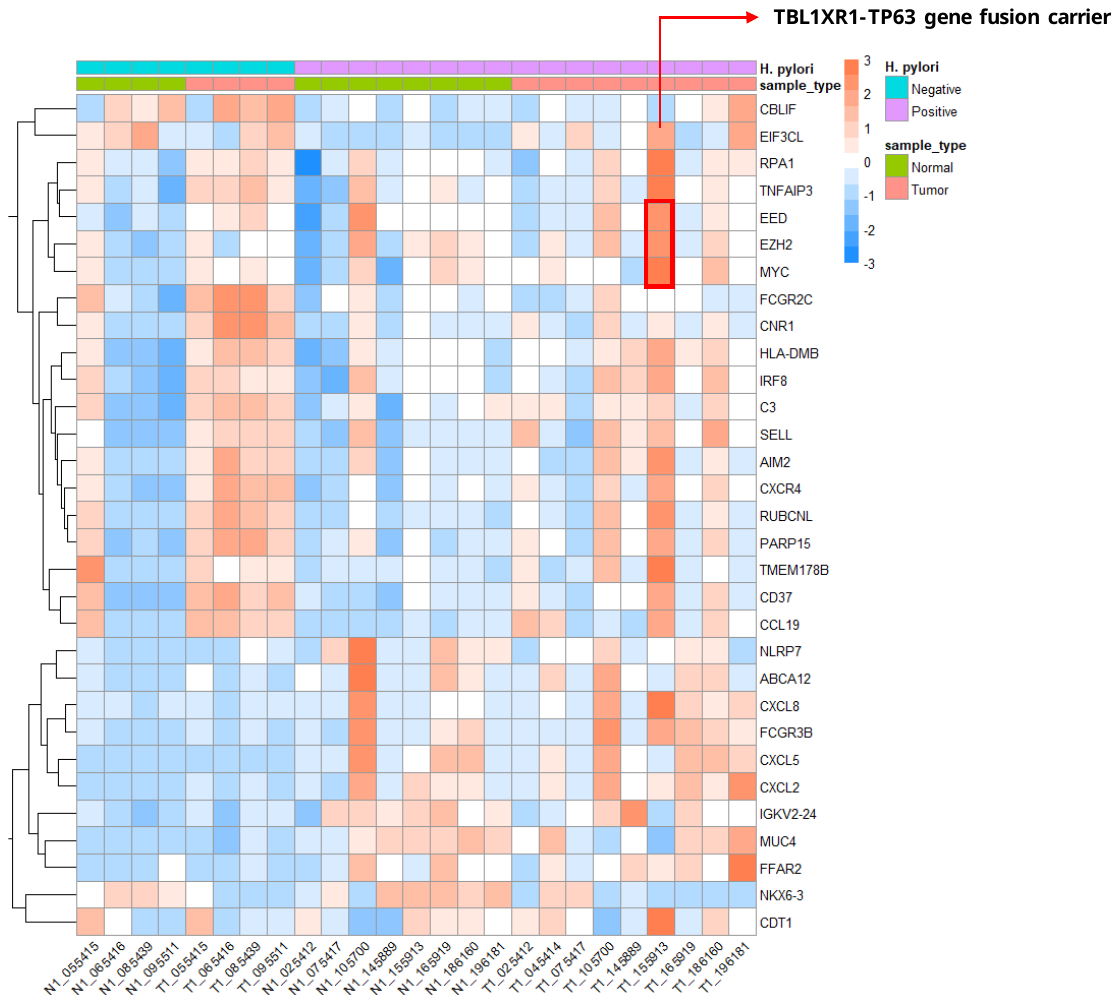
**Figure S1.** A heatmap shows the relative expression levels of genes which were statistically significant in stepwise comparison analyses or manually selected.
